# Structure-based modeling of SARS-CoV-2 peptide/HLA-A02 antigens

**DOI:** 10.1101/2020.03.23.004176

**Authors:** Santrupti Nerli, Nikolaos G. Sgourakis

## Abstract

As a first step toward the development of diagnostic and therapeutic tools to fight the Coronavirus disease (COVID-19), it is important to characterize CD8+ T cell epitopes in the SARS-CoV-2 peptidome that can trigger adaptive immune responses. Here, we use RosettaMHC, a comparative modeling approach which leverages existing high-resolution X-ray structures from peptide/MHC complexes available in the Protein Data Bank, to derive physically realistic 3D models for high-affinity SARS-CoV-2 epitopes. We outline an application of our method to model 439 9mer and 279 10mer predicted epitopes displayed by the common allele HLA-A*02:01, and we make our models publicly available through an online database (https://rosettamhc.chemistry.ucsc.edu). As more detailed studies on antigen-specific T cell recognition become available, RosettaMHC models of antigens from different strains and HLA alleles can be used as a basis to understand the link between peptide/HLA complex structure and surface chemistry with immunogenicity, in the context of SARS-CoV-2 infection.

---

An ongoing pandemic caused by the novel SARS coronavirus (SARS-CoV-2) has become the focus of extensive efforts to develop vaccines and antiviral therapies (1). Immune modulatory interferons, which promote a widespread antiviral reaction in infected cells, and inhibition of pro-inflammatory cytokine function through anti-IL-6/IL-6R antibodies, have been proposed as possible COVID-19 therapies (2, 3). However, stimulating a targeted T cell response against specific viral antigens is hampered by a lack of detailed knowledge of the immunodominant epitopes displayed by common Human Leukocyte Antigen (HLA) alleles across individuals (public epitopes). The molecules of the class I major histocompatibility complex (MHC-I, or HLA in humans) display on the cell surface a diverse pool of 8 to 15 amino acid peptides derived from the endogenous processing of proteins expressed inside the cell (4). This MHC-I restriction of peptide antigens provides jawed vertebrates with an essential mechanism for adaptive immunity: surveillance of the displayed peptide/MHC-I (pMHC-I) molecules by CD8+ cytotoxic T-lymphocytes allows detection of aberrant protein expression patterns, which signify viral infection and can trigger an adaptive immune response (5). A recent study has shown important changes in T cell compartments during the acute phase of SARS-CoV-2 infection (6), suggesting that the ability to quantify antigen-specific T cells would provide new avenues for understanding the expansion and contraction of the TCR repertoire in different disease cohorts and clinical settings. Given the reduction in breadth and functionality of the naïve T cell repertoire during aging (7), identifying a minimal set of viral antigens that can elicit a protective response will enable the design of diagnostic tools to monitor critical gaps in the T cell repertoire of high-risk cohorts, which can be addressed using peptide or epitope string DNA vaccines (8).

Human MHC-I molecules are extremely polymorphic, with thousands of known alleles in the classical HLA-A, -B and -C loci. Specific amino acid polymorphisms along the peptide-binding groove (termed A-F pockets) define a repertoire of 10^4^-10^6^ peptide antigens that can be recognized by each HLA allotype (9, 10). Several machine-learning methods have been developed to predict the likelihood that a target peptide will bind to a given allele (reviewed in (11)). Generally these methods make use of available data sets in the Immune Epitope Database (12) to train artificial neural networks that predict peptide processing, binding and display, and their performance varies depending on peptide length and HLA allele representation in the database. Structure-based approaches have also been proposed to model the bound peptide conformation *de novo* (reviewed in (13)). These approaches utilize various algorithms to optimize the backbone and side chain degrees of freedom of the peptide/MHC structure according to an all-atom scoring function, derived from physical principles (14–16), that can be further enhanced using modified scoring terms (17) or mean field theory (18). While these methods do not rely on large training data sets, their performance is affected by bottlenecks in sampling of different backbone conformations, and any possible structural adaptations of the HLA peptide-binding groove.

Predicting the bound peptide conformation whose N- and C-termini are anchored within a fixed-length groove is a tractable modeling problem that can be addressed using standard comparative modeling approaches (19). In previous work focusing on the HLA-B*15:01 and HLA-A*01:01 alleles in the context of neuroblastoma neoantigens, we have found that a combined backbone and side chain optimization approach can yield accurate pMHC-I models for a pool of target peptides, provided that a reliable template of the same allele and peptide length can be identified in the database (20). In this approach (RosettaMHC), a local optimization of the backbone degrees of freedom is sufficient to capture minor (within 0.5 Å heavy atom RMSD) changes of the target peptide backbone relative to the conformation of the peptide in the template, used as a starting point. For HLA-A*02:01, the most common HLA allele among disease-relevant population cohorts (21), there is a large number of high-resolution X-ray structures available in the PDB (22), suggesting that a similar principle can be applied to produce models of candidate epitopes directly from the proteome of a pathogen of interest. Here, we apply RosettaMHC to all HLA-A*02:01 epitopes predicted directly from the ~30 kbp SARS-CoV-2 genome, and make our models publicly available through an online database. The computed binding energies of our models can be used as an additional validation layer to select high-affinity epitopes from large peptide sets. As detailed epitope mapping data from high-throughput tetramer staining (23–25) and T cell functional screens (26) become available, the models presented here can provide a toehold for understanding links between pMHC-I antigen structure and immunogenicity, with actionable value for the development of peptide vaccines to combat the disease.

## Materials and Methods

### Identification of SARS-CoV-2 peptide epitopes

The SARS-CoV-2 protein sequences (https://www.ncbi.nlm.nih.gov/nuccore/NC_045512.2) were obtained from NCBI and used to generate all possible peptides of lengths 9 and 10 (9,621 9mer and 9,611 10mer peptides). We used NetMHCpan-4.0 (27) to derive binding scores to HLA-A*02:01, and retained only peptides classified as strong or weak binders (selected using the default percentile rank cut-off values). The binding classification was performed using eluted ligand likelihood predictions. While in this study we use NetMHCpan-4.0 predictions as inputs to select candidate epitopes for structure modeling, our workflow is fully compatible with any alternative epitope prediction method.

### Selection of PDB templates

To model SARS-CoV-2 / HLA-A*02:01 antigens, we identified 3D structures from the PDB that can be used as templates for comparative modeling. First, we selected all HLA-A02 X-ray structures that are below 3.5 Å resolution and retained only those that have 100% identity to the HLA-A*02:01 heavy chain sequence (residues 1-180). We found 241 template structures bound to epitopes of lengths from 8 to 15 residues (of which 170 are 9mers and 61 are 10mers). For each SARS-CoV-2 target peptide, we selected a set of candidate templates of the same length by matching the target peptide anchor positions (P2 and P9/P10) to each peptide in the template structures. Then, we used the BLOSUM62 (28) substitution matrix to score all remaining positions in the pairwise alignment of the target/template sequences, and the PDB template with the top alignment score was selected for modeling. For target peptides where we found no templates which match both peptide anchors, we scored all positions in the pairwise alignment and selected the top scoring template for modeling.

### RosettaMHC modeling framework and database

RosettaMHC (manuscript in preparation) is a comparative modeling protocol developed using PyRosetta (29) to model pMHC-I complexes. The program accepts as input a list of peptide sequences, an HLA allele definition and a template PDB file (selected as described in the previous step). To minimize “noise” in the simulation from parts of the MHC-I fold that do not contribute to peptide binding, only the α_1_ and α_2_ domains are considered in all steps. For each peptide, a full alignment between the target and template peptide/MHC sequences is performed using clustal omega (30). The alignment is used as input to Rosetta’s threading protocol (*partial_thread.<ext>*). From the threaded model, all residues in the MHC-I groove that are within a heavy-atom distance of 3.5 Å from the peptide are subjected to 10 independent all-atom refinement simulations using the FastRelax method (31) and a custom movemap file. Binding energies are extracted from the refined structures using interface analyzer protocol (*InterfaceAnalyzer.<ext>*). The top three models are selected based on the binding energies, and used to compute an average energy for each peptide in the input list. RosettaMHC models of SARS-CoV-2/HLA-A*02:01 epitopes are made available through an online database (see data availability). The website that hosts our database was constructed using the Django web framework.

## Results and Discussion

### Template identification for structure modeling using RosettaMHC

Our full workflow for template identification and structure modeling is outlined in Figure 1a, with a flowchart shown in Figure 1b. To identify all possible regular peptide binders to HLA-A*02:01 that are expressed by SARS-CoV-2, we used a recently annotated version of all open reading frames (ORFs) in the viral genome from NCBI (32), made available through the UCSC genome browser (33). We used 9- and 10-residue sliding windows to scan all protein sequences, since these are the optimum peptide lengths for binding to the HLA-A*02:01 groove (34). While spliced peptide epitopes (35) are not considered in the current study, this functionality can be added to our method at a later stage. Using NetMHCpan-4.0 (27), we identified all 439 9mer and 279 10mer epitopes that are predicted to yield positive (classified as both weak and strong) binders. To further validate this set and derive plausible 3D models of the peptide/HLA-A*02:01 complexes, we used a structure-guided approach, RosettaMHC, which aims to derive a physically realistic fitness score for each peptide in the HLA-A*02:01 binding groove using an annotated database of high-resolution structures and Rosetta’s all-atom energy function (36). RosettaMHC leverages a database of 241 HLA-A*02:01 X-ray structures encompassing a range of bound peptides, to find the closest match to each target epitope predicted from the SARS-CoV-2 proteome. To identify the best template for structure modeling, we use sequence matching criteria which first consider the peptide anchors (positions P2 and P9/P10 for 9mer/10mer epitopes), followed by a sequence similarity metric calculated from the full alignment between the template and target peptide sequences. The template assignment statistics for the four different classes of SARS-CoV-2 epitopes in our set are shown in Figure 2a. We find that we can cover the entire set of 718 predicted binders using a subset of 114 HLA-A*02:01 templates in our annotated database of PDB-derived structures (Figure 2b). Each target peptide sequence is then threaded onto the backbone of its best identified template, followed by all-atom refinement of the side chain and backbone degrees of freedom using Rosetta’s Ref2015 energy function (36), and binding energy calculation.

**FIGURE 1.**
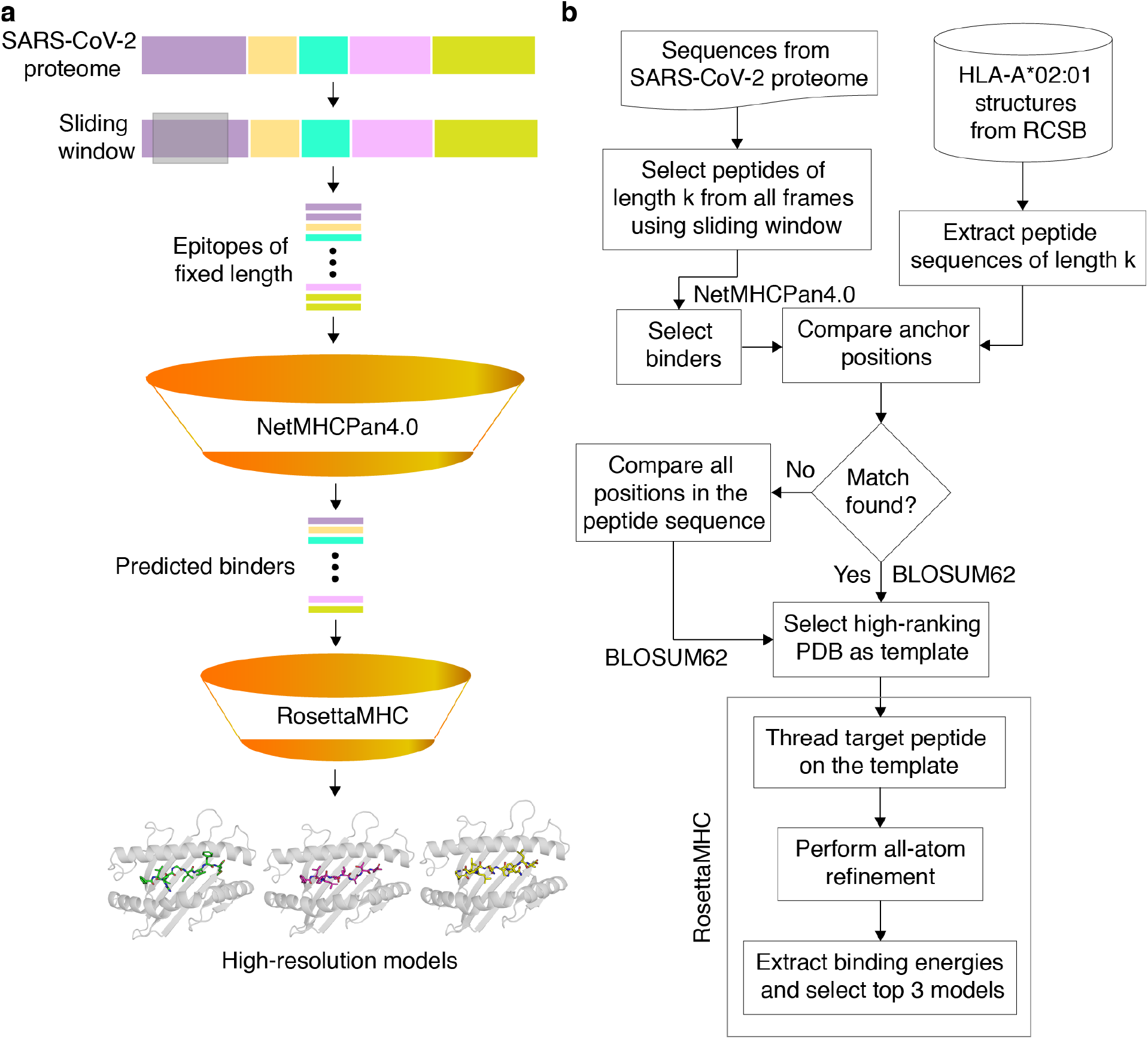
Structure-guided modeling of T cell epitopes in the SARS-CoV-2 proteome. **(a)** General workflow of our pipeline for structure-guided epitope ranking. **(b)** Protein sequences from the annotated SARS-CoV-2 proteome are used to generate peptide epitopes with a sliding window covering all frames of a fixed length (9,621 9mer and 9,611 10mer possible peptides). Candidate peptides are first filtered by NetMHCpan-4.0 (27) to identify all predicted strong and weak binders (439 9mer and 279 10mer epitopes). For rapid template matching and structure modeling, we use a local database of 241 HLA-A*02:01 X-ray structures with resolution below 3.5 Å from the Protein Data Bank (22). Each candidate peptide is scanned against all peptide sequences of the same length in the database, and the top-scoring template is used to guide the RosettaMHC comparative modeling protocol and to compute a binding energy.

**FIGURE 2.**
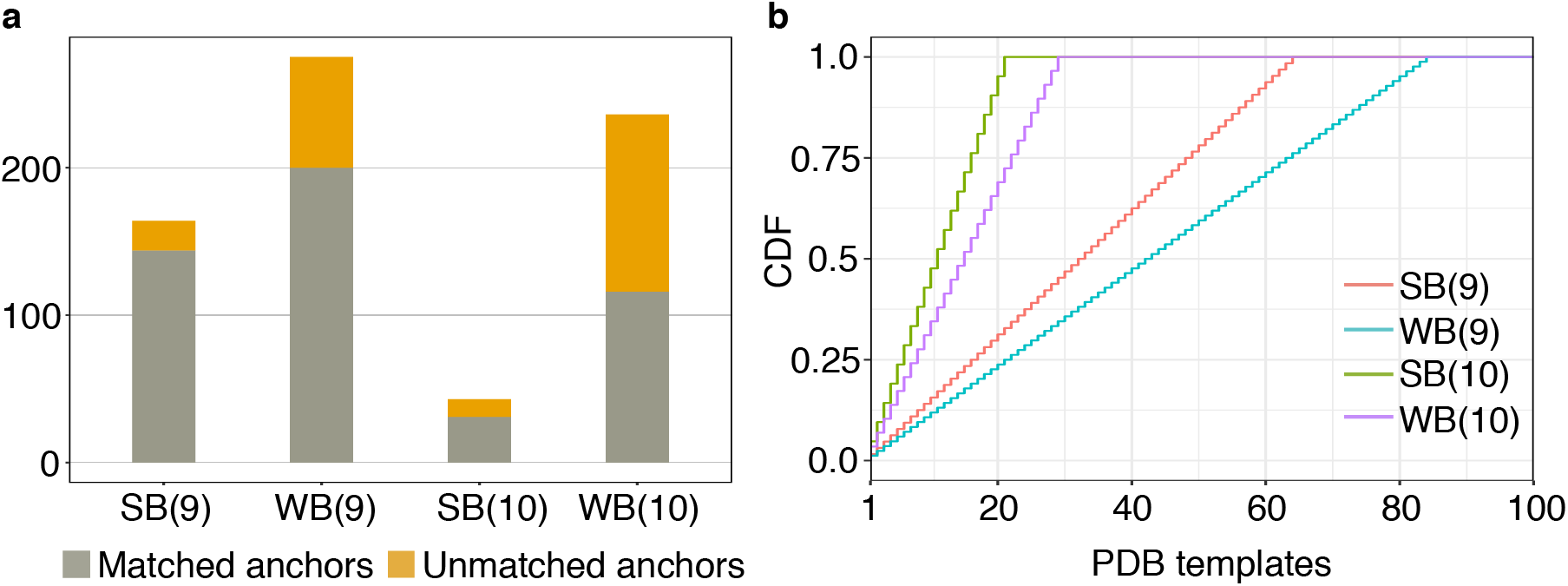
Coverage of predicted HLA-A02 epitopes by structural templates in the PDB. **(a)** Peptide anchor matching statistics of all predicted SARS-CoV-2 strong (SB) and weak binders (WB) of lengths 9 and 10 to a database of 241 high-resolution HLA-A*02:01 X-ray structures **(b)** Plot showing cumulative distribution (CDF) of strong and weak binder peptides of lengths 9 and 10, as a function of the total number of matching templates from the Protein Data Bank (22).

### RosettaMHC models recapitulate features of high-resolution X-ray structures

The sequence logos derived from 9mer and 10mer peptides with good structural complementarity to the HLA-A*02:01 groove according to Rosetta’s binding energy (see below) adhere to the canonical motif, with a preference for hydrophobic, methyl-bearing side chains at the peptide anchor residues P2 and P9 (Figure 3a). The anchor residue preferences are recapitulated in representative 9mer and 10mer models of the two top binders in our set as ranked by Rosetta’s energy (Figure 3c and 3d), corresponding to epitopes TMADLVYAL and FLFVAAIFYL derived from the RNA polymerase and nsp3 proteins, respectively, which are both encoded by *orf1ab* in the viral genome (NCBI Reference YP_009724389.1). In accordance with features seen in high-resolution structures of HLA-A*02:01-restricted epitopes, the peptides adopt an extended, bulged backbone conformation. The free N-terminus of both peptides is stabilized by a network of polar contacts with Tyr 7, Tyr 159, Tyr 171 and Glu 63 in the A- and B- pockets of the HLA-A*02:01 groove. The Met (9mer) or Leu (10mer) side chain of P2 is buried in a B-pocket hydrophobic cleft formed by Met 45 and Val 67. Equivalently, the C-terminus is coordinated through polar contacts with Asp 77 and Lys 145 from opposite sides of the groove, with the Leu P9/P10 anchor nestled in the F-pocket defined by the side chains of Leu 81, Tyr 116, Tyr 123 and Trp 147. Residues P3-P8 form a series of backbone and side chain contacts with pockets C, D and E, while most backbone amide and carbonyl groups form hydrogen bonds with the side chains of residues lining the MHC-I groove. These high-resolution structural features are consistent across low-energy models of unrelated target peptides in our input set, suggesting that, when provided with a large set of input templates, a combined threading and side chain optimization protocol can derive physically realistic models.

**FIGURE 3.**
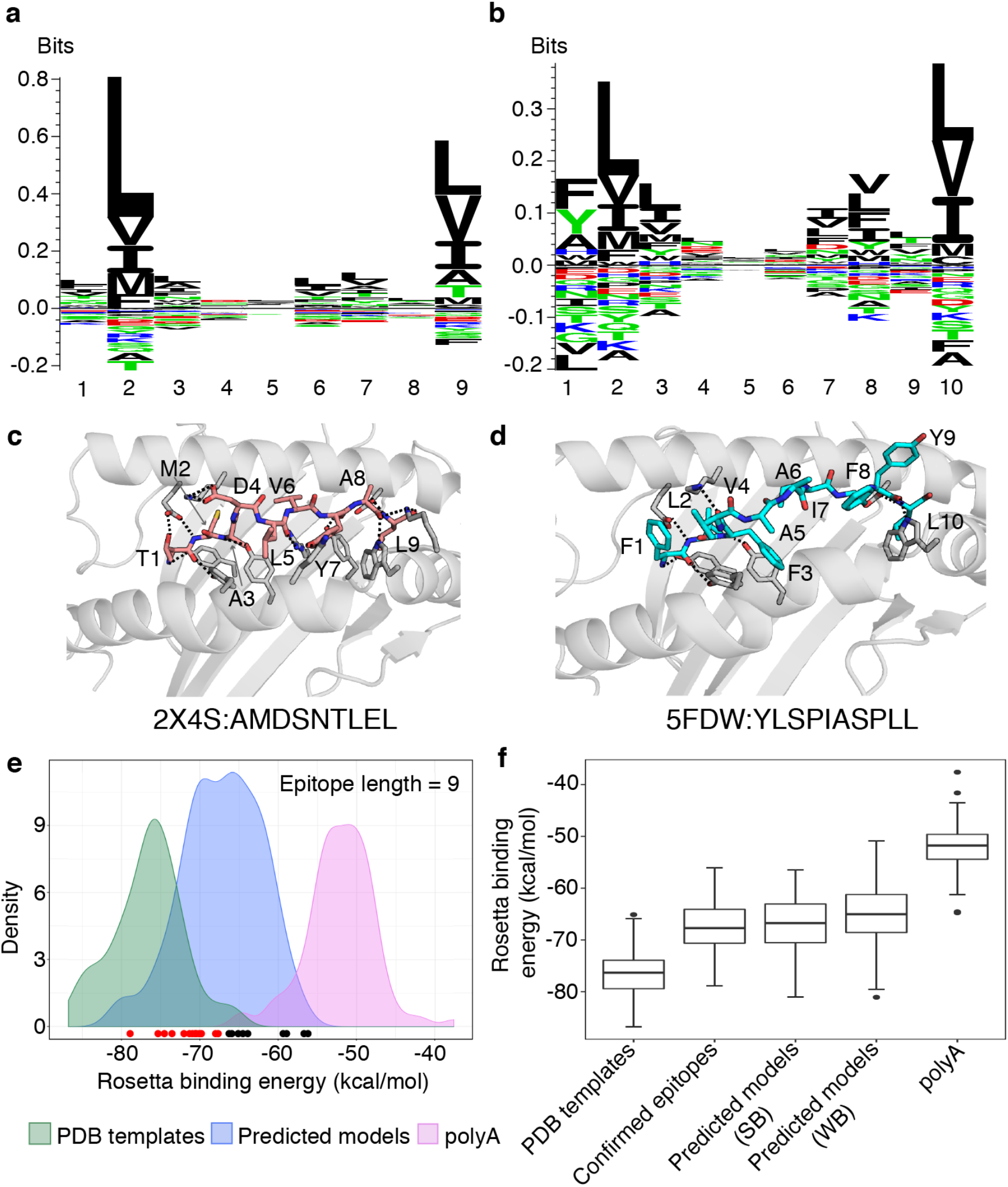
Summary of RosettaMHC modeling results for SARS-CoV-2 peptide epitopes. Sequence logos from the *n* top ranking epitopes in the SARS-CoV-2 genome, predicted by NetMHCpan-4.0 (27) and further refined using RosettaMHC binding simulations are shown for: **(a)** 9mers (*n*=154) and **(b)** 10mers (*n*=72). The top 9mer and 10mer epitopes in our refined set are shown: **(c)** TMADLVYAL, from RNA polymerase and **(d)** FLFVAAIFYL, from nsp3. Dotted lines indicate polar contacts between peptide and heavy chain residues, with peptide residues labelled. The template PDB IDs and original peptides used for modeling the target peptides are indicated below each model. **(e)** Density plots showing distribution of average Rosetta binding energies (kcal/mol) for all epitopes of length 9. Distributions reflect 93 PDB templates (green), 164 strong binder epitopes (according to NetMHCpan-4.0 (27)) (blue), and 93 poly alanine peptides modeled using the same PDB templates and used as a reference set for sub-optimal binders (polyA; pink). The binding energies of models generated for 28 confirmed SARS T cell epitopes from the IEDB and ViPR (37, 38) are indicated by circles at the bottom of the plot. Red circles (19/28) indicate epitopes that lie within the distribution of refined PDB templates and black circles (9/28) indicate epitopes that fall within the distribution of polyA (sub-optimal binders). **(f)** Box plots showing distribution of average binding energies for 93 PDB templates, 93 poly alanine peptides, 28 confirmed epitopes (37, 38) and RosettaMHC models for 164 strong (SB) and 275 weak (WB) binder 9mer epitopes predicted from the SARS-CoV-2 proteome using NetMHCpan-4.0 (27).

### Selection of high-affinity peptide epitopes using a structure-based score

To evaluate the accuracy of our models and fitness of each peptide within the HLA-A*02:01 binding groove, we computed Rosetta all-atom binding energies across all complexes modeled for different peptide sets. High binding energies can be used as an additional metric to filter low-affinity peptides in the NetMHCpan-4.0 predictions, with the caveat that high energies can be also due to incomplete optimization of the Rosetta energy function as a result of significant deviations between the target and template backbone conformations, not captured by our protocol. We performed 10 independent calculations for each peptide, and the 3 lower-energy models were selected as the final ensemble and used to compute an average binding energy. The results for all 9mer peptides are summarized in Figures 3e, f, while additional results for 10mers are provided through our web-interface and outlined in Supplemental Table 1. As a positive reference, we used the binding energies of the idealized and relaxed PDB templates, which are at a local minimum of the Rosetta scoring function. As a reference set for sub-optimal binders, we modeled decoy structures of poly alanine (polyA) peptide sequences (predicted by NetMHCpan-4.0 to be a top 9th percentile binder for HLA-A*02:01), threaded onto the same PDB templates.

We observe a significant, negative (−26 kcal/mol) energy gap between the average binding energies for PDB templates and poly alanine models. The binding energies for all modeled 9mers from the SARS-CoV-2 genome fall between the average energies of the optimal PDB templates and sub-optimal polyA binders, and show a bimodal distribution with significant overlap with the refined PDB template energies (Figure 1e). Comparison of the distributions between epitopes that are classified as strong versus weak binders by NetMHCpan-4.0 shows a moderate bias towards lower binding energies for the strong binders and a larger spread in energies for weak binders, likely due to suboptimal residues at the P2 and P9 anchor positions (Figure 3f). As an intendent positive set, we also modeled 28 9mer peptides that are homologous to peptides in the SARS viral genome and have been previously reported to bind HLA-A*02:01 in the IEDB and ViPR (12, 37, 38) databases (Supplemental Table 2). Inspection of Rosetta binding energies derived from models in this set shows a similar distribution to the epitopes classified by NetMHCpan-4.0 as strong binders, with the energies of 19/28 peptides falling well within the distribution of the refined PDB templates (red dots in Figure 3e).

Based on these observations, we further classified all epitopes in the original set provided by NetMHCpan-4.0 as strong or weak binders according to the Rosetta binding energy. Peptides with binding energies that fall well within the PDB template distribution (green curve and red dots in Figure 3e) are classified as strong binders. We obtained 154 9mer and 72 10mer strong binders which show optimal complementarity within the HLA-A*02:01 peptide-binding groove according to our modeling simulations. These results suggest that the high-resolution features seen in our models (Figure 3c, d) yield optimal binding energies for a significant fraction of the epitopes predicted by NetMHCpan-4.0 (45/33% of strong binders and 30/25% of weak binders for 9mers/10mers, respectively), which are comparable to locally refined PDB structures. The average binding energies for all peptides are provided in our web-interface and in Supplemental Table 1.

### Surface features of peptide/HLA-A*02:01 models for T cell recognition

Visualization of our models through an interactive online interface provides direct information on SARS-CoV-2 peptide residues that are bulging out of the MHC-I groove, and are therefore accessible to interactions with complementarity-determining regions (CDRs) of T cell receptors (TCRs). Given that αβ TCRs generally employ a diagonal binding mode to engage pMHC-I antigens where the CDR3α and CDR3β TCR loops form direct contacts with key peptide residues (39, 40), knowledge of the surface features for different epitopes adds an extra layer of information to interpret sequence variability between different viral strains. For other important antigens with known structures in the PDB, such features can be derived from an annotated database connecting pMHC-I/TCR co-crystal structures with biophysical binding data (41), and were recently employed in an artificial neural network approach to predict the immunogenicity of different HLA-A*02:01 bound peptides in the context of tumor neoantigen display (42). A separate study has shown that the electrostatic compatibility between self vs foreign HLA surfaces can be used to determine antibody alloimmune responses (43). Given that antibodies and TCRs use a common fold and similar principles to engage pMHC-I molecules (40), it is likely that surface electrostatic features play an important role in recognition of peptide/HLA surfaces by their cognate TCRs in the context of SARS-CoV-2 infection.

Electrostatic surface potentials calculated using a numerical solution to the Poisson-Boltzmann Equation (44) for our modeled peptide/HLA-A*02:01 complexes allow us to compare important features for TCR recognition between different high-affinity epitopes (Figure 4). We observe a moderate electropositive character of the HLA-A*02:01 α_1_ helix, and a moderate negative potential on the α2 helix, which is consistent between complexes with different bound peptides. However, due to substantial sequence variability in surface-exposed residues at the P2-P8 positions, we observe a range of electrostatic features ranging from negative (epitope TMADLVYAL), to neutral (NLIDSYFVV) or positively charged (KLWAQCVQL). Further classification and ranking of the top binders in our set on the basis of their molecular surface features would enable the selection of the most diverse panel of peptides for high-throughput pMHC tetramer library generation (23–25). Tetramer screening of T cells from COVID-19 patients, recovered individuals and healthy donors can be used to identify critical gaps in the T cell repertoire of high-risk groups, and to design epitope DNA strings for vaccine development.

**FIGURE 4.**
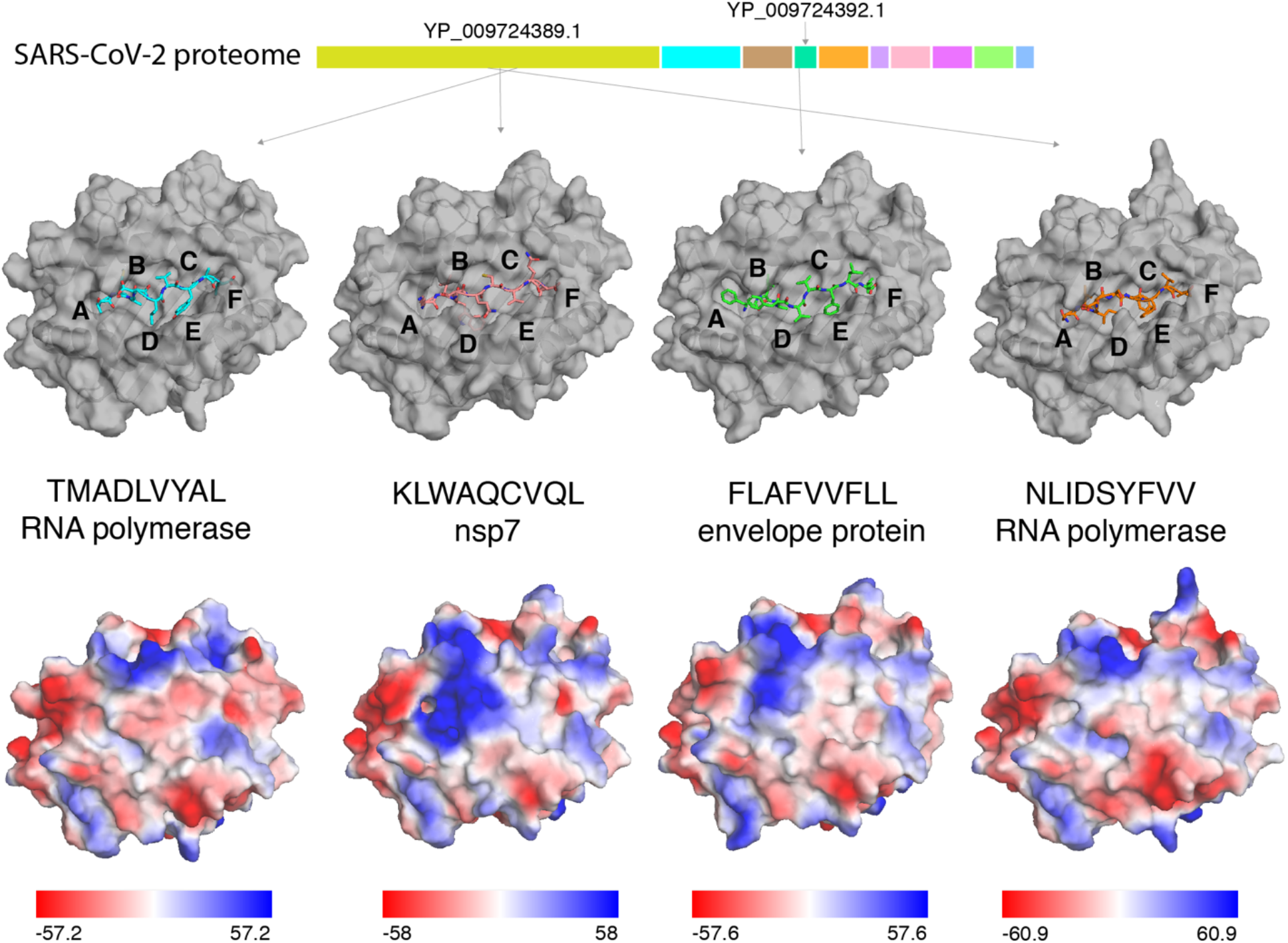
Variability in TCR recognition features of HLA-A02 with different high-affinity peptides. Molecular surfaces of SARS-CoV-2/HLA-A*02:01 RosettaMHC models are shown for four top-scoring epitopes (ranked by Rosetta binding energy from left to right) captured in the A, B, C, D, E and F pockets of the MHC-I groove (top panel). The origins of the peptide epitopes in the ~30 kbp SARS-CoV-2 genome are noted. Electrostatic surfaces computed for the same models are shown in the bottom panel. Solvent-accessible surface representation with electrostatic potential in the indicated ranges (down to −60 kcal/(mol·*e*) in red and up to +61 kcal/(mol·*e*) in blue) were calculated using the APBS solver (45) in Pymol (46). All calculations were performed at 150 mM ionic strength, 298.15 Kelvin, pH 7.2, protein dielectric 2.0, and solvent dielectric 78.54. Electrostatic potentials are given in units of kT/e. A 1.4 Å solvent (probe) radius and 10.0 points/Å^2^ density was used to calculate molecular surfaces.

## Acknowledgements

The authors are grateful to Alison Lindberg and ITS/ADC staff at UCSC for assistance in setting up the web server for the database, and Hiram Clawson (UCSC Genome Browser) for providing the SARS-CoV-2 protein sequence data. We thank Andrew McShan, Hailey Wallace (UCSC) for assistance in the development of the RosettaMHC protocol, and David Haussler (UCSC), Michael Betts (University of Pennsylvania) and David Margulies (NIH) for helpful discussions. This work was supported through Grants from NIAID (5R01AI143997) and NIGMS (5R35GM125034).

## Code and Data availability

An online web-interface for visualization and download of all models is available at: https://rosettamhc.chemistry.ucsc.edu. The RosettaMHC source code is available at https://github.com/snerligit/mhc-pep-threader. Rosetta binding energies for all 718 HLA-A*02:01-restricted peptides in our set are provided in Supplemental Table 1.

